# Scale-free structure of cancer networks and their vulnerability to hub-directed combination therapy

**DOI:** 10.1101/2020.07.01.159657

**Authors:** Andrew X. Chen, Christopher J. Zopf, Jerome Mettetal, Wen Chyi Shyu, Joseph Bolen, Arijit Chakravarty, Santhosh Palani

## Abstract

**Background:** The effectiveness of many targeted therapies is limited by toxicity and the rise of drug resistance. A growing appreciation of the inherent redundancies of cancer signaling has led to a rise in the number of combination therapies under development, but a better understanding of the overall cancer network topology would provide a conceptual framework for choosing effective combination partners. In this work, we explore the scale-free nature of cancer protein-protein interaction networks in 14 indications. Scale-free networks, characterized by a power-law degree distribution, are known to be resilient to random attack on their nodes, yet vulnerable to directed attacks on their hubs (their most highly connected nodes).

**Results:** Consistent with the properties of scale-free networks, we find that lethal genes are associated with ∼5-fold higher protein connectivity partners than non-lethal genes. This provides a biological rationale for a hub-centered combination attack. Our simulations show that combinations targeting hubs can efficiently disrupt 50% of network integrity by inhibiting less than 1% of the connected proteins, whereas a random attack can require inhibition of more than 30% of the connected proteins.

**Conclusions:** We find that the scale-free nature of cancer networks makes them vulnerable to focused attack on their highly connected protein hubs. Thus, we propose a new strategy for designing combination therapies by targeting hubs in cancer networks that are not associated with relevant toxicity networks.

## Background

Despite the many successes against cancer in the laboratory, progress in the clinic has been slow. Today, as was true a generation ago, cancer is primarily treated with surgery, radiation and combination chemotherapies, which are effective at the early stages of cancer development, but typically show limited success in increasing the long-term survival of late-stage cancer patients (Brenner *et al*., 2009). Recently, the focus has shifted towards targeted agents that seek to inhibit specific molecular signaling pathways used by cancer cells for growth, cell division, and metastasis (Sawyers, 2004; Bozic *et al*., 2012). By shutting down signaling pathways, targeted agents seek to disrupt the protein-protein interaction networks (signaling pathways) that provide function and structure to cancer cells. However, apart from a few high-profile successes (*e*.*g*. Gleevec, Zelboraf) that specifically target mutant proteins produced by cancers, targeted therapy still remains ineffective due to systemic toxicity that precludes dosing these agents at high levels (Lee, 2012), as well as due to the adaptive nature of cancer cells that exploit the redundancy in their complex signaling networks (Gillies *et al*., 2012; Gyurkó *et al*., 2013; Peterson *et al*., 2012; Thompson *et al*., 2015).

Features of network complexity such as crosstalk and feedback can hinder treatment efficacy, such as with EGFR (Yamaguchi *et al*., 2014) or mTOR kinase inhibitors (Rodrik-Outmezguine *et al*., 2011). Recent studies have highlighted the inverse correlation between cancer patient survival and both the complexity (Breitkreutz *et al*., 2012) and modularity (Takemoto *et al*., 2013) of molecular signaling networks. In this context, a better understanding of cancer network complexity may be valuable in the design of new therapies (Hopkins, 2007; Jacunski *et al*., 2013; Pe’er *et al*., 2011). A number of studies have demonstrated the utility of network theory in cancer for drug discovery (Durmuş *et al*., 2009; Dejori *et al*., 2004), classification (Chuang *et al*., 2007), and prognosis (Li *et al*., 2010; Taylor *et al*., 2009; Ergün *et al*., 2007). A growing appreciation of the complexity and inherent redundancy of cancer cell signaling networks has spurred renewed interest in the development of new combination therapies (Komarova *et al*., 2009; Bozic *et al*., 2013). However, the selection of combination agents for specific cancer indications lacks a unifying framework that exploits the underlying architecture and vulnerabilities present in these complex networks.

In this work, we seek to apply findings from graph theory to develop a conceptual framework for the selection of effective combination partners. Graph theory provides a basis for connecting network topology to functional properties. Graph theory has revealed that many real-world networks such as airline hubs and the Internet are scale-free (Barabasi *et al*., 1999). Scale-free networks are structured as highly connected hub nodes surrounded by a large number of poorly connected spokes. Since the majority of nodes are only weakly connected, such networks are robust to randomly selected node removals (Cohen *et al*., 2000). On the other hand, the highly connected hubs form an integral part of the network, leaving them vulnerable to targeted attacks (Cohen *et al*., 2001). Translating this graph theory notion to cancer networks, cancer-associated proteins and their interactions are mapped to nodes and their edges (Ben-Tal *et al*., 2014). Previous studies have suggested scale-free network architectures in glioblastoma (Ladha *et al*., 2010), gastric cancer (Aggarwal *et al*., 2006), and colon cancer (Ruan *et al*., 2006) though these results are based partially or primarily on correlations in gene expression, a surrogate for network edges. Highly connected proteins are known to be enriched for genes associated with cancer (Schramm *et al*., 2010). If cancer networks are indeed scale-free, this graph theory insight encourages the intuitive notion that these highly connected proteins are also more vital for the cancer cell’s survival (Jeong *et al*., 2001; Yu *et al*., 2004; He *et al*., 2006; Albert 2005). By successively inhibiting proteins of cancer networks with the most interactions, the overall structure of the network may be destroyed – thus eliminating network sources of resistance such as redundancy, crosstalk or feedback. This hub-directed strategy may therefore provide an approach to increase clinical efficacy.

In this work, we show that cancer networks are likely scale-free for 14 different indications based on protein-protein signaling and connectivity data. We demonstrate the biological significance of targeting highly connected protein hubs in cancer networks by showing that lethal genes have higher numbers of protein-protein interactions relative to non-lethal genes. In simulating multiple hub knockouts, we find that combination therapy results in a rapid and steady decrease in cancer network integrity in all 14 indications tested. Therefore, extensive damage to cancer cells may only require inhibiting a small fraction of the cancer-associated proteins. Taking account of the vulnerabilities of scale-free cancer networks, we propose a framework that successively removes the highest connected nodes of the network that are not shared with pertinent toxicity networks as a design for targeted combination therapy.

## Methods

### Scale Free Nature of Cancer Networks

To construct the protein-protein network for each cancer indication, we obtained cancer-associated proteins for 14 available indications from the KEGG PATHWAY database (Kanehisa *et al*., 2014; Kanehisa *et al*., 2000) and identified their interactions using the BioGRID human gene interaction repository, Version 3.2.113 (Stark *et al*., 2006). Each KEGG pathway was mapped to the subset of the repository of interactions including at least one of the pathway’s proteins. This generated an indication-specific network comprising interactions related to the pathway. Only unique, physical, non-self interactions, such as those acquired by two-hybrid or affinity apture ethods were selected. This list of interactions was imported into Cytoscape (Shannon *et al*., 2003), a network visualization software, to create a schematic of the cancer network (Figs. 1A and S1A). The degree (number of interactions) of each protein was counted in MATLAB (The Mathworks Inc., Natick, MA), generating a degree distribution for each network. The MATLAB package *plfit* (Clauset *et al*., 2009), which uses maximum-likelihood methods to bin and truncate data, was used to estimate the power-law parameters and plot the fit (Figs. 1B and S1B). An exponential fit, using the same bins from *plfit* was also determined (Fig. S1A). R2 values were calculated for each power law fit by linear regression of the log-log plot, and for each exponential fit by linear regression of the log plot. Additionally, a log-likelihood ratio (as per equation C.3 in Clauset *et al*., 2009) was used to compare the probability of the power law fit versus the exponential fit. A schematic of this workflow is visualized in Fig. S5.

**Figure 1.**
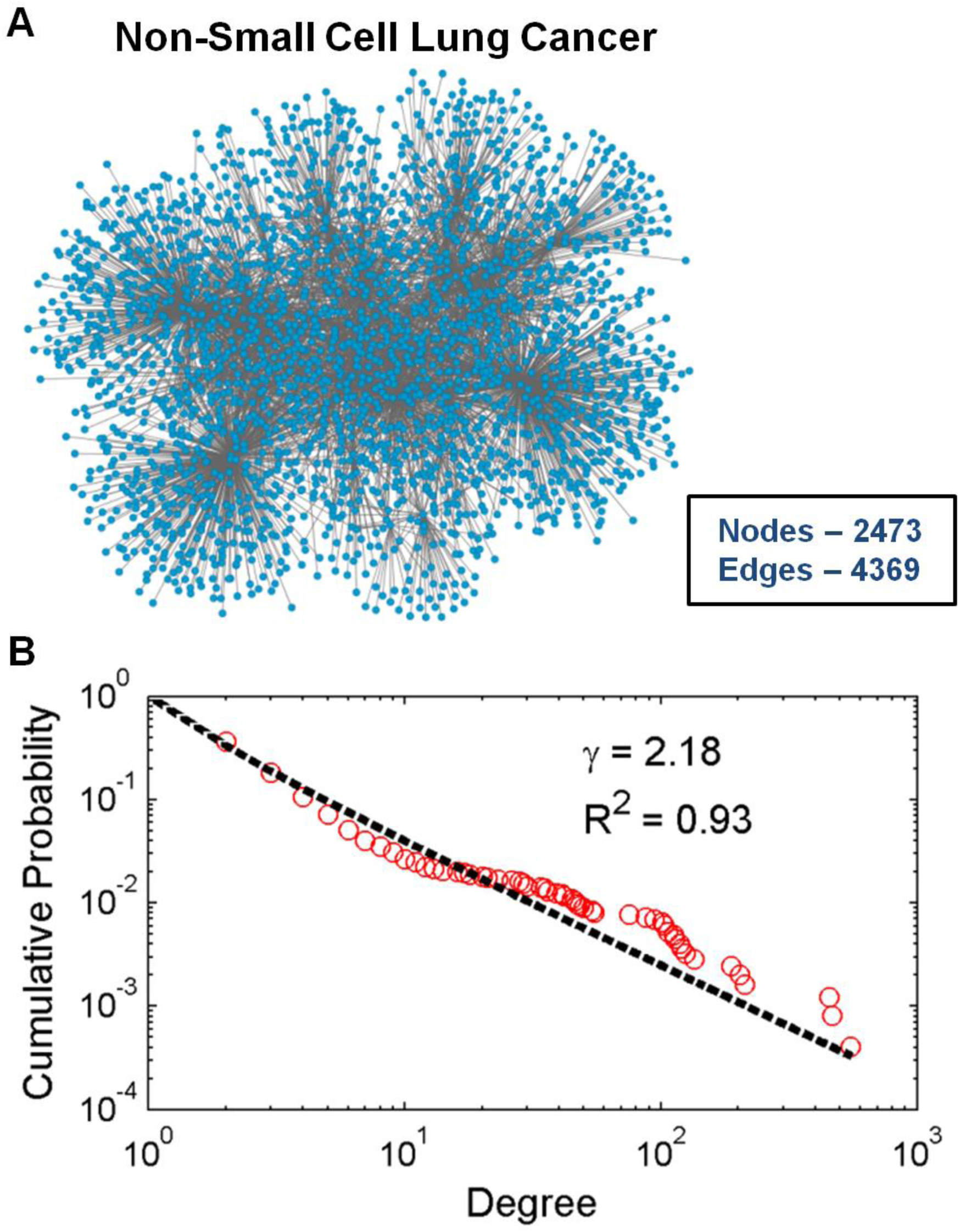
Cancer networks exhibit scale-free characteristics. (A) Visualization of protein-protein interactions of the genes associated with the non-small cell lung cancer indication as described in the Methods. (B) The cumulative degree distribution (red circles) for the network in (A) is fitted to a power law function (black dotted lines) with the coefficient γ shown.

### Connectivity-Lethality Correlation

To determine whether highly connected proteins tended to also be more lethal, we cross-referenced protein-protein interaction data from BioGRID with genome-wide RNA interference data from a liquid tumor cell line (Tiedemann *et al*., 2012) and a solid tumor cell line (Sethi *et al*., 2012). These RNA interference studies of multiple myeloma and ovarian cancer cells presented viability data of druggable genomes consisting of 6722 and 6022 genes, and identified 160 and 300 lethal hits, respectively. Secondary confirmation studies were then used to validate 57 and 53 lethal targets, of which 51 and 52 were found within BioGRID. The druggable genomes were cross-referenced to the BioGRID database to find all physical interactions where at least one gene was part of the druggable genome. This interaction list was imported into Cytoscape to visualize the network structure and to demonstrate its scale-free nature through a power-law fit (Figs. S2A and S2B).

To compare the degree-connectivity of the lethal genes to the full network, we computed their probability distributions. The positions of the 51 lethal genes in the multiple myeloma network and 52 lethal genes in the ovarian cancer network were noted within the degree-connectivity ranking, and used to generate a cumulative distribution function (CDF) for the probability that a given node was lethal, similar to a lethality curve (Jeong *et al*., 2003). This CDF was fit to a nonlinear Hill function, generating estimates for the median degree-connectivity of lethal genes compared to all genes. Using the fitted CDFs, the difference in median connectivity between lethal and all genes was divided by the standard error of the fit found for the lethal CDF, leading to a z-score and corresponding p-value.

### Citability-Lethality Correlation

To verify that the connectivity-lethality correlation is not a result of citation bias, we next tested whether or not the enrichment observed could be explained by a citability-lethality correlation. The number of citations for each gene was determined from the NCBI Gene database (Maglott *et al*., 2005). Only genes that were part of the druggable genome were used. Akin to the connectivity analysis, the citation distributions of lethal genes from ovarian cancer and multiple myeloma were compared to the entire networks. The cumulative citation frequency was calculated for both cancers, and the median citation-enrichment was observed.

### Analysis of Network Collapse

To quantify a measure of network disintegration, we investigated the behavior of the giant component in response to successive hub removal. The giant component is defined as the largest spanning cluster size, a global metric of network integrity (Cohen *et al*., 2002). The analysis of giant component reduction was done in accordance with the methods described in (Cohen *et al*., 2001). In brief: the size of the largest spanning cluster was estimated as the most-connected nodes were successively removed. This value was normalized to the size of the original spanning cluster. In analyzing networks under random attacks, simulations removing randomly selected nodes were used to assess the average node removals needed to attain 50% giant component reduction.

### Distance-Giant Component Correlation

To investigate the impact of inter-hub distance on network destruction, 10,000 scale-free networks were randomly generated by the Barabasi-Albert model (Barabasi *et al*., 1999) with N = 1000 and m = 1. This corresponds to 1 node being added at a time to a network with size similar to the 14 cancer networks. The seed network was 5 nodes connected in a line. The highest and second highest-degree nodes were identified, and their closest distance computed by Dijkstra’s algorithm (Dijkstra *et al*., 1959). These two nodes were then removed from the network (along with all their edges), and the sizes of the largest remaining components were calculated (Fig. S4A). The giant component reduction was averaged after collating networks by their closest hub distance.

## Results

To determine whether cancer networks are scale-free, we examined the interaction networks for 14 indications constructed from the KEGG cancer pathways and BioGRID database. These networks’ sizes and properties are listed in Table 1. Visualization of the networks (Fig. 1A, Fig S1A) revealed a large extent of centrality, with nodes that appeared to be structured as hubs and spokes. Each node is assigned a degree for its total number of interactions, which yields a degree distribution representing all of the network’s nodes (Fig. 1B, Fig S1B).

**Table 1.** Properties of cancer networks. For 14 cancer indications, the number of nodes (proteins) and edges (interactions) are listed for that indication’s network. The cumulative degree distribution of these networks was fitted to a power law (Power) and an exponential (Exp), resulting in a power law coefficient (gamma), and R2 of the fits. Additionally, the log-likelihood of the power law fit was compared to the exponential fit. The number and percentages of nodes required to reduce the network’s giant component (GC) to 50% is listed for hub-directed and randomly selected node removal strategies.

| Network | Nodes | Edges | Gamma | Fit R<sup>2</sup> |  | Log-Likelihood Ratio | Hub Removals for 50% GC Reduction |  | Random Node Removals for 50% GC Reduction |  |
| --- | --- | --- | --- | --- | --- | --- | --- | --- | --- | --- |
|  |  |  |  | Power | Exp |  | # | % | # | % |
| Colorectal cancer | 2906 | 5356 | 2.37 | 0.92 | 0.63 | $2.3 \cdot 10^3$ | 23 | 0.79 | 1040 | 35.8 |
| Pancreatic cancer | 3158 | 6113 | 2.31 | 0.92 | 0.64 | $2.5 \cdot 10^3$ | 29 | 0.92 | 1033 | 32.7 |
| Glioma | 3133 | 5655 | 2.17 | 0.92 | 0.58 | $8.1 \cdot 10^3$ | 19 | 0.61 | 1101 | 35.1 |
| Thyroid cancer | 1664 | 2459 | 2.43 | 0.89 | 0.55 | $4.6 \cdot 10^3$ | 7 | 0.42 | 565 | 34.0 |
| Acute myeloid leukemia | 2475 | 4441 | 2.18 | 0.94 | 0.61 | $5.7 \cdot 10^3$ | 22 | 0.89 | 805 | 32.5 |
| Chronic myeloid leukemia | 3737 | 8149 | 2.26 | 0.92 | 0.63 | $3.4 \cdot 10^3$ | 37 | 0.99 | 1408 | 37.7 |
| Basal cell carcinoma | 1224 | 1577 | 2.68 | 0.85 | 0.55 | $3.7 \cdot 10^3$ | 4 | 0.33 | 425 | 34.7 |
| Melanoma | 2553 | 4360 | 2.23 | 0.91 | 0.57 | $6.8 \cdot 10^3$ | 15 | 0.59 | 894 | 35.0 |
| Renal cell carcinoma | 2667 | 4920 | 2.4 | 0.92 | 0.62 | $2.2 \cdot 10^3$ | 22 | 0.82 | 909 | 34.1 |
| Bladder cancer | 2518 | 4113 | 2.29 | 0.88 | 0.56 | $7.1 \cdot 10^3$ | 10 | 0.40 | 809 | 32.1 |
| Prostate cancer | 4114 | 8908 | 2.24 | 0.93 | 0.62 | $3.9 \cdot 10^3$ | 36 | 0.88 | 1465 | 35.6 |
| Endometrial cancer | 2733 | 4732 | 2.23 | 0.91 | 0.58 | $7.0 \cdot 10^3$ | 16 | 0.59 | 896 | 32.8 |
| Small cell lung cancer | 3319 | 6815 | 2.3 | 0.93 | 0.61 | $3.2 \cdot 10^3$ | 33 | 0.99 | 1239 | 37.3 |
| Non-small cell lung cancer | 2473 | 4369 | 2.18 | 0.93 | 0.58 | $6.0 \cdot 10^3$ | 19 | 0.77 | 882 | 35.7 |

The defining characteristic of a scale-free network is a power-law decrease of its degree distribution as compared to exponential tails of randomly constructed networks. When the cumulative degree distribution of these networks was fit to a power-law (Fig. 1B, Fig. S1B), the R^2^was generally around 0.9, and provided a better description of the distribution than an alternative exponential model (R^2^∼ 0.6) (Table 1). Additionally, log-likelihoods ratios of over 10^3^ favor power law over exponential fits for each indication. In particular, the exponential fit cannot describe the presence of high-connection nodes – an unlikely occurrence in a randomly constructed network. The power-law coefficients of these fits ranged from 2.1 to 2.7, which are typical for realistic scale-free networks (Barabasi *et al*., 1999) that have degree distributions with well-defined means.

Scale-free network properties imply the existence of highly vulnerable and highly connected hub proteins. To understand the biological significance of hub-directed attack in a scale-free cancer network, we cross-referenced the degree-connectivity of proteins with their lethality from genome-wide siRNA knockout screens performed on multiple myeloma (Tiedemann *et al*., 2012) and ovarian cancer cells (Sethi *et al*., 2012). By comparing the degree distribution of only the validated lethal hits to the whole druggable genome, we observed an overall positive shift in the cumulative degree distribution of the lethal genes compared to all genes in the network for both the cancer indications (Fig. 2). Furthermore, the median connectivity of lethal genes for the ovarian cancer and multiple myeloma networks are 6-fold and 4-fold higher than the average gene median connectivity, respectively. Comparing the cumulative distribution fits, we observed a statistically significant (*p-values* < 10^−6^) enrichment in connectivity in lethal genes compared to all druggable genes.

**Figure 2.**
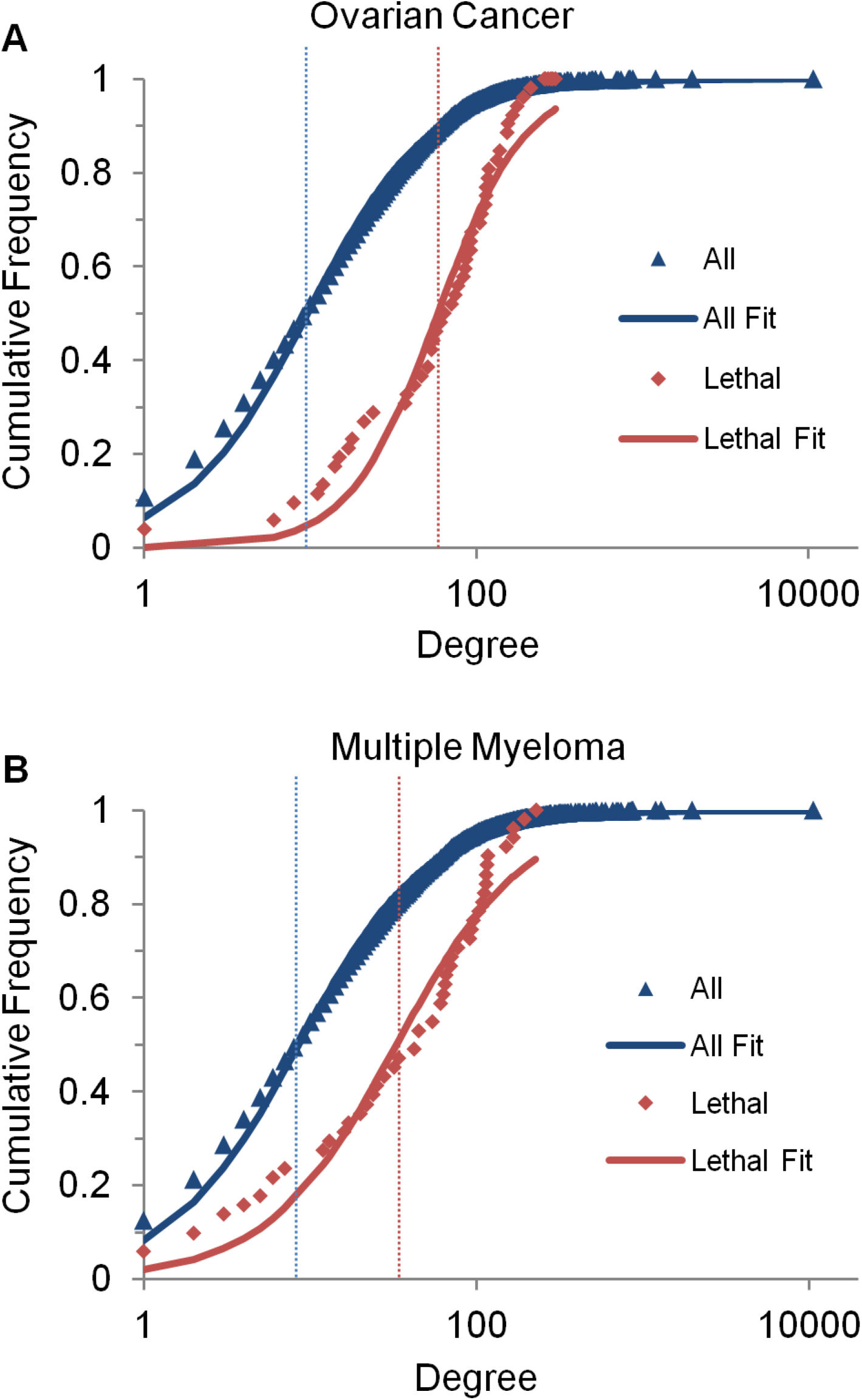
Lethal genes have a higher degree-connectivity than the average gene. Degree-connectivity for all druggable and lethal genes in both ovarian cancer (A) and multiple myeloma (B) cell lines were assigned their degree-connectivity within the network as described in Methods. Hill function fits of the degree-connectivity (thick lines) provide estimates of median degree (vertical lines) displaying approximately 6-fold and 4-fold enrichments for the lethal genes in the ovarian and multiple myeloma networks, respectively (p-values of 10^−36^ and 10^−6^, respectively).

Based on the results that cancer networks are scale-free and lethal genes are more likely to be hubs, we simulated a combination therapy strategy of sequentially targeting cancer protein hubs. Understanding that combined sets of node removals may cause more network damage than their individual knockouts, we utilized giant component reduction (signifying the change in largest spanning cluster size, a global metric of network integrity) to quantify the extent of network damage. For a successively increasing number of hub knockouts, we simulated the giant component reduction of each of the 14 cancer networks (Fig. 3A). We found that cancer networks are efficiently disintegrated by the sequential removal of their most highly connected nodes. Surprisingly, the basal cell carcinoma network is nearly destroyed with the knockout of only the top 10 hubs, and all networks attained 50% reduction of their giant component with removal of less than 1% of the full network nodes (a maximum of 37 hub proteins). In contrast, when randomly selected nodes were sequentially removed, 50% giant component reduction typically required removal of over 30% of the network’s nodes (Table 1). Furthermore, we observe that the marginal utility of destroying each successive hub is roughly linear for most networks (Fig. 3B). A linear marginal utility implies that this strategy does not saturate and will lead to greater network destruction with every additional hub removal. Altogether, our findings show that combinations of hub-targeted drugs can efficiently reduce the cancer network’s integrity.

**Figure 3.**
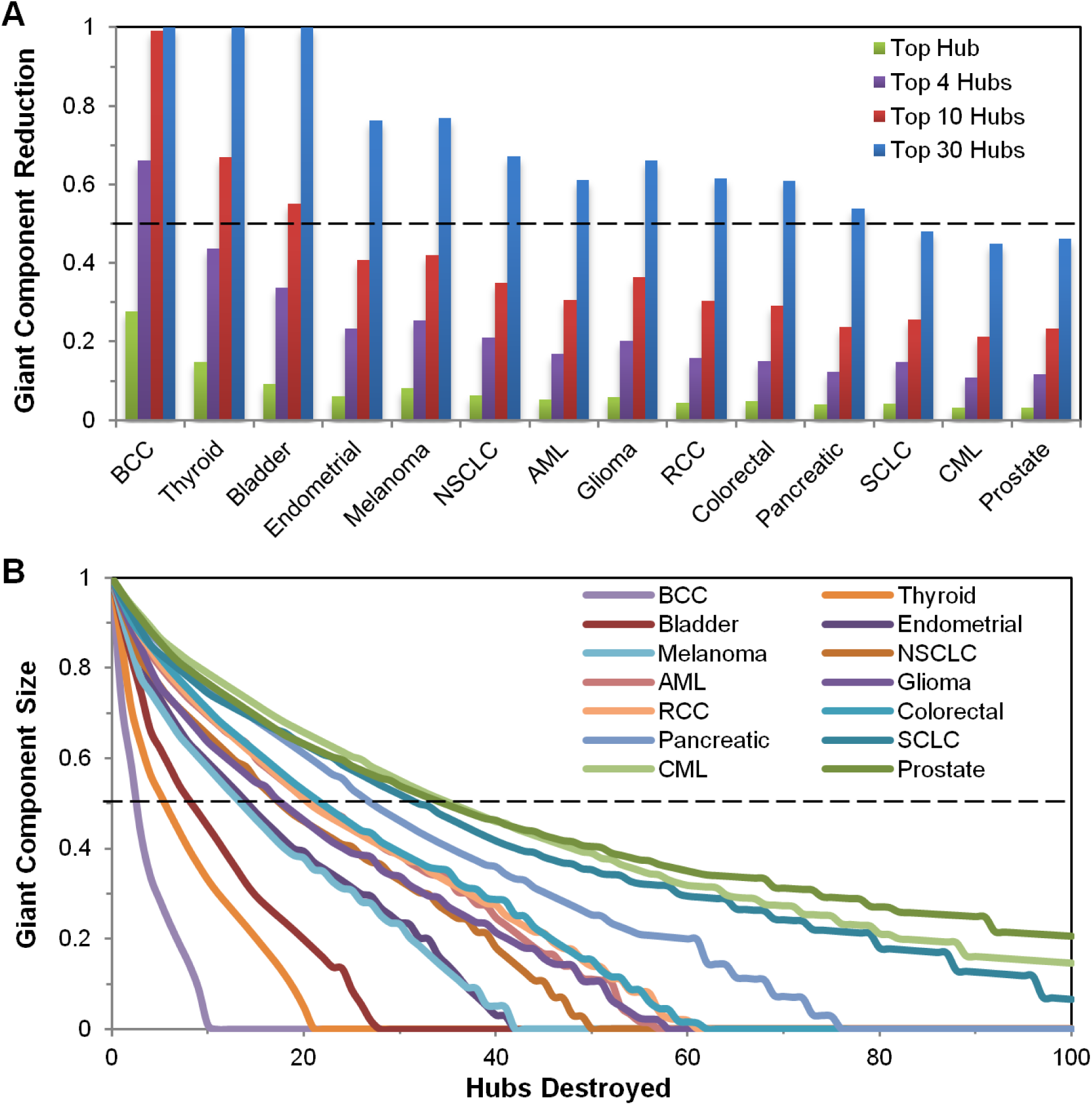
Successive knockouts of cancer network hubs quickly destroy network integrity. (A) The giant component reduction of 14 cancer networks (with their 1^st^ neighbors) was calculated as the highest-connected hubs (Top 1, 4, 10 and 30) were successively removed. (B) Reduction in the giant component size is shown as a function of number of hubs destroyed. A linear marginal utility in giant component reduction is seen across all indications with a hub-directed attack.

## Discussion

Cancer is a complex disease that relies on pathway redundancies in biological networks to evolve and confer resistance to drugs (Peterson *et al*., 2012). Drug combinations are highly effective if they target a disease that relies on few genes for its growth and development. An example is HIV, the life cycle of which is primarily dependent on 9 genes that govern cell-entry, reverse transcription, host DNA integration and virion production. By developing drugs that target each of the phases of its life cycle and administering them in combination, multiple obstacles prevent HIV replication in spite of frequently ongoing gene mutation (Oversteegen *et al*., 2007). One can imagine using a similar brute-force approach to treat cancer; however, blocking all the functional pathways contributing to cancer cell viability will require an infeasible order of magnitude of combined drugs. On the other hand, if we understand the underlying architecture of the signaling pathway, we can potentially identify vulnerabilities that will allow us to use a realistic combination scale to combat cancer.

In this work, we have demonstrated that cancer protein-protein networks are much more likely to be scale-free than random. While there is controversy about how well-fit a power law must be (Khanin *et al*., 2006), it is clear that these networks possess a few highly connected proteins that contribute largely to the integrity of the overall network. This is supported by our finding that lethal genes tend to have higher protein connectivity partners (4- to 6-fold) than non-lethal genes in cancer networks, thereby providing a biological rationale for a hub-directed attack with combination therapy. Since lethal proteins can be potential drug targets, it is possible that they are better studied and hence have more documented protein-protein interactions. To account for this confounding effect, we constructed CDFs of citation counts for the druggable genome and lethal genes. The enrichment in the citation count seen in lethal genes was modest (1.3 – 1.4-fold) relative to the druggable genome (Fig. S3), suggesting that citation bias does not explain the observed enrichment in degree count for lethal genes.

The redundancies built into cancer signaling networks may require the destruction of a large number of proteins to definitively eliminate output from a functional pathway. Our results suggest that cancer networks can be efficiently destroyed by the simultaneous removal of a few highly connected hubs. Even though there are thousands of proteins associated with each indication, we demonstrate that the removal of less than 1% of proteins is sufficient to create 50% reduction in the integrity of all cancer networks investigated. In practice, the required amount of cluster reduction, and therefore the number of drugs in a combination, may be even smaller. While the level of giant component reduction required for a significant decrease in viability is hard to pinpoint, these results demonstrate broad network disruption can be achieved by eliminating a very small fraction of the overall cancer network. Additionally, as knowledge of the druggable genome continues to expand, the proposed strategy for identifying highly connected, lethal targets will become more and more tractable.

One of the primary limitations to combination drug therapy is the emergence of toxicity. The efficacy network in Fig. 4 represents a schematic of the protein-protein interactions associated with disease (e.g. breast cancer), while the toxicity network denotes protein-protein connections related with normal function (e.g. neutrophil production). Most research on developing cancer therapies has focused primarily on constructing efficacy networks for specific indications to identify drug targets. The approach presented here allows us to characterize the efficacy protein networks to identify the critical proteins (hubs 1, 2 and 3 in Fig. 4) for combination therapy. However, hub proteins are likely to be connected with proteins with diverse functionalities, so targeting some of the hubs (hub 2 in Fig. 4) can lead to unacceptable toxicity (e.g. high grade neutropenia) as it interferes with normal function. Hence, understanding the protein-protein interactions of relevant toxicity networks is a crucial step in successfully implementing the hub-directed combination approach. Assembling and characterizing the protein networks of most common dose-limiting toxicities will enable us to enhance the therapeutic window by targeting only the efficacy-specific hubs and avoiding the toxicity-inducing hubs.

**Figure 4.**
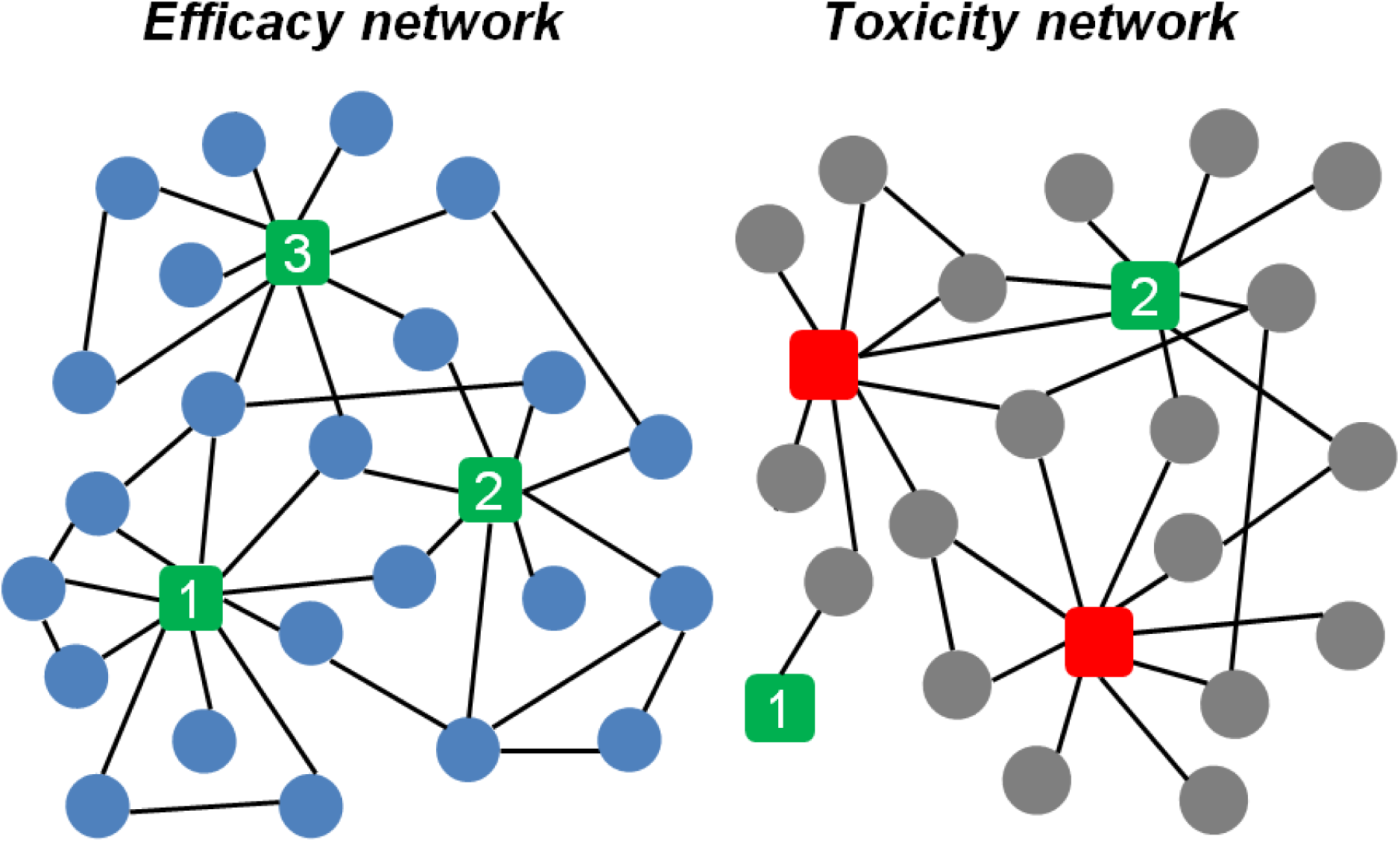
Differences between efficacy and toxicity networks may be exploitable. Low-connectivity nodes in the efficacy and toxicity network are shown as solid blue and grey circles, respectively, and hubs are represented as squares. Hubs 1 and 3 are efficacy-specific hubs while hub 2 is shared between the efficacy and toxicity networks. Red squares represent toxicity-specific hubs.

Here we have focused only on the degree-connectivity of the network in identifying vulnerabilities, as it provides a direct correlation to lethality. However, there may be more to destroying a network than targeting based on node degree alone (Kovacs *et al*., 2015). For example, a hub-directed attack on a cancer network could be more effective if it leverages the relative context of the hubs to one another in the network (Rachlin *et al*., 2006). Supporting the notion that local topology may be important, we found through simulations that in scale-free networks generated with the Barabási–Albert model the shorter the distance between the removed hubs, the larger the reduction in network integrity (Supp Fig. 4). Hence, in addition to a hub-directed approach, exploiting local hub topologies may provide further ammunition to fully capitalize on the vulnerabilities of the cancer networks.

## Conclusions

Our findings provide a conceptual framework to design drug combinations by targeting the hubs present in the cancer networks while avoiding the hubs present in relevant toxicity networks. Viewed more generally, the work provides a justification for focusing on highly connected hub proteins as a basis for designing combination therapies against the highly interconnected protein networks of cancer.

## Availability of Data and Material

The datasets supporting the conclusions of this article are available online at the Kyoto Encyclopedia of Genes and Genomes (http://www.genome.jp/kegg/) and the BioGRID (http://thebiogrid.org/).

## Acknowledgements

We would like to thank Eric Lightcap for discussions on combination therapy, as well as the DMPK department of Takeda Boston for their support.

