## Supplementary Figures for "Scale-free structure of cancer networks and their vulnerability to hub-directed combination therapy"

**Figure S1. KEGG cancer networks follow power law degree distributions. (A)** As described in Figure 1, the KEGG genes associated with 14 cancer pathways were referenced using the BioGRID database to generate interaction networks. All 14 KEGG networks are visualized, displaying their scale-free architecture. **(B)** These networks were well-described by power law distributions (black lines) with coefficient γ between 2.1 and 2.7. When fitted to exponential distributions (blue lines), the $R^2$ was lower across all indications.

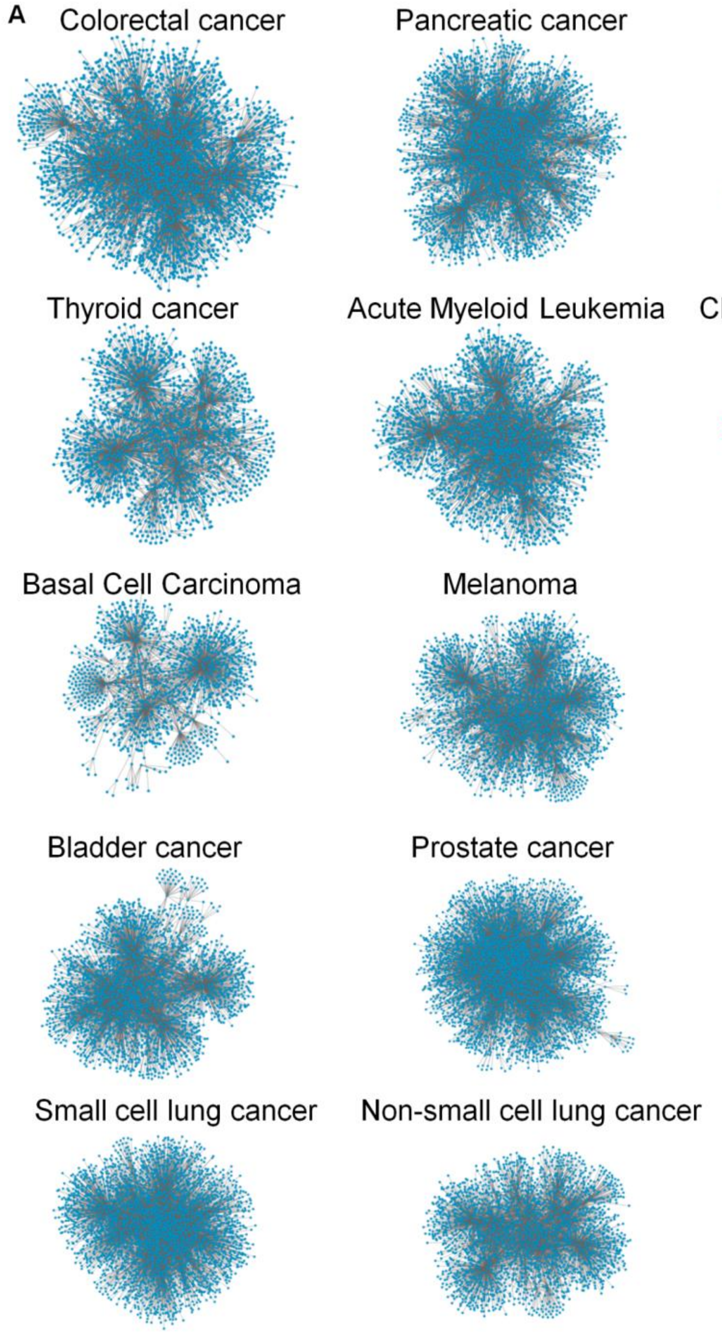

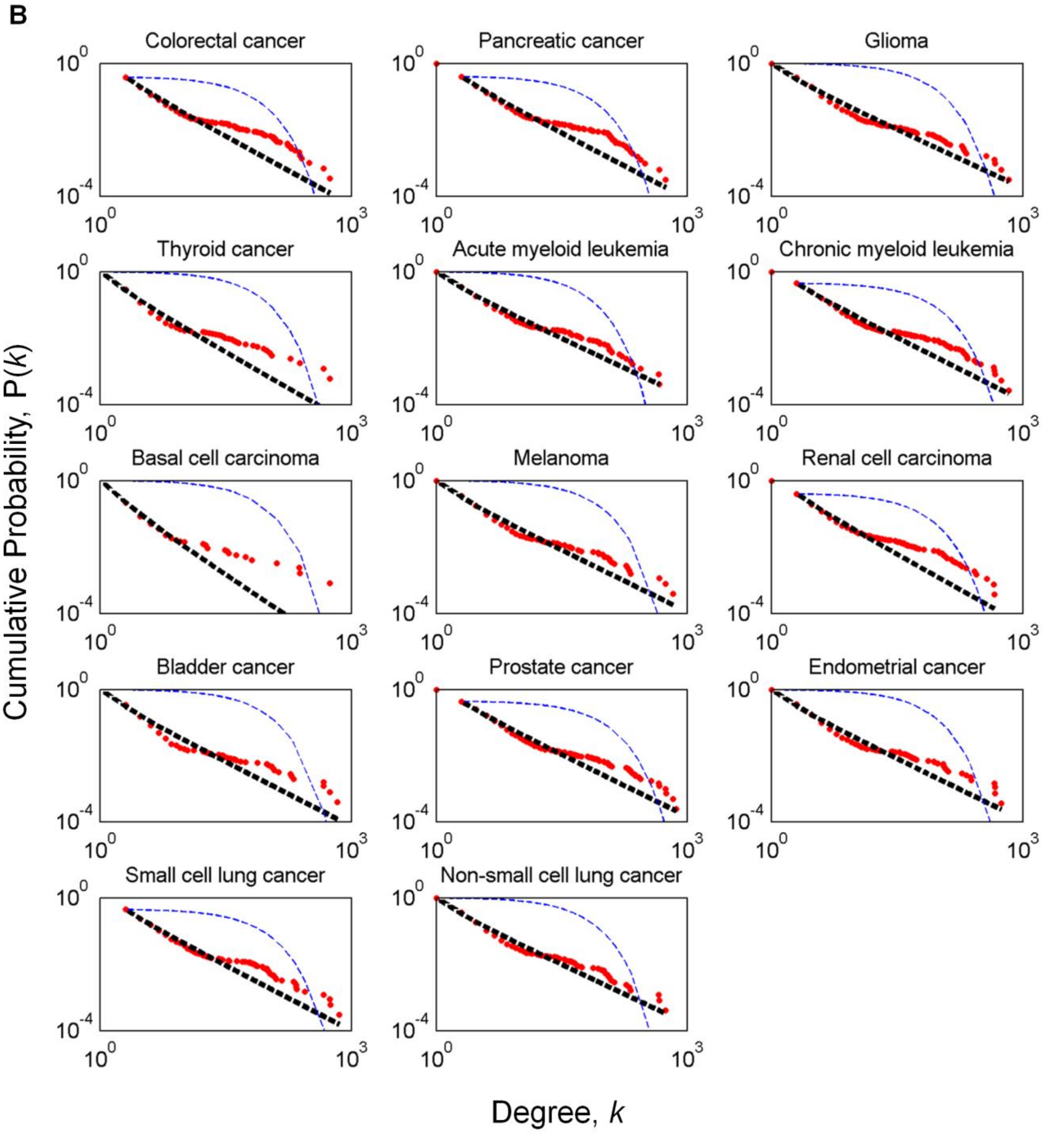
B
Colorectal cancer
Pancreatic cancer
Glioma
Thyroid cancer
Acute myeloid leukemia
Chronic myeloid leukemia
Basal cell carcinoma
Melanoma
Renal cell carcinoma
Bladder cancer
Prostate cancer
Endometrial cancer
Small cell lung cancer
Non-small cell lung cancer
$10^0$
$10^{-4}$
$10^3$
Cumulative Probability, P(*k*)
Degree, *k*

**Figure S2. BioGRID networks exhibit scale-free characteristics.** Visualization of protein-protein interactions of the genes associated with ovarian cancer (A) and multiple myeloma (B). The protein nodes list was obtained from druggable genomes listed in Sethi (2012) and Tiedemann (2012), and the physical protein-protein interaction details were captured from the BioGRID human dataset. (C, D) The degree distribution (red dots) for the networks above is fitted to a power law function (black dotted lines) to check for scale-freedom.

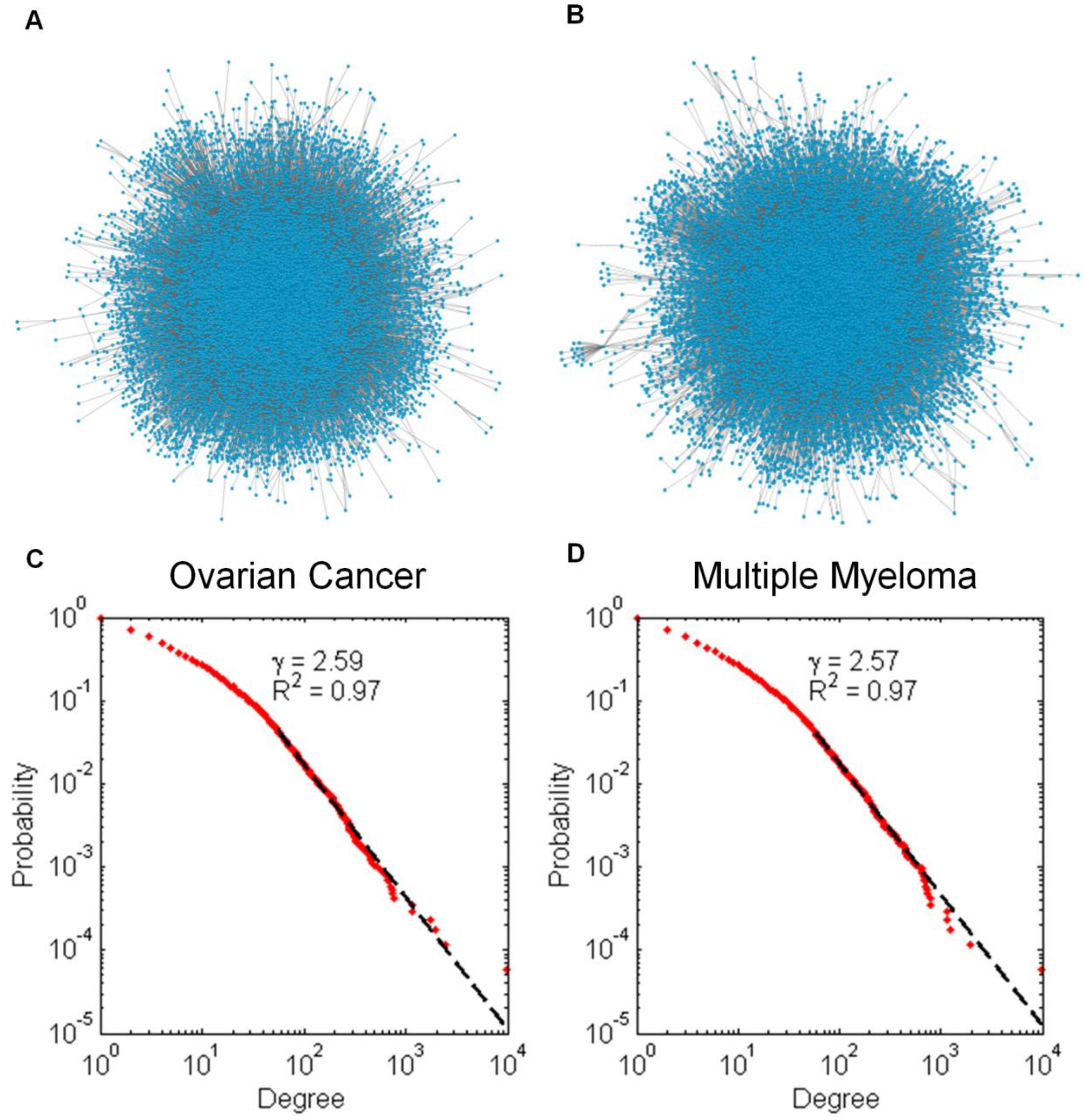

**Figure S3. Correlation of Lethal Genes with Citation Count.** The cumulative distribution of nodes in respect to their citation count for ovarian cancer (A) and multiple myeloma (B) shows that lethal genes are somewhat enriched. For multiple myeloma and ovarian cancer, the median degrees are enriched by factors of 1.3 and 1.4, corresponding to p-values of $10^{-55}$ and $10^{-188}$, respectively.

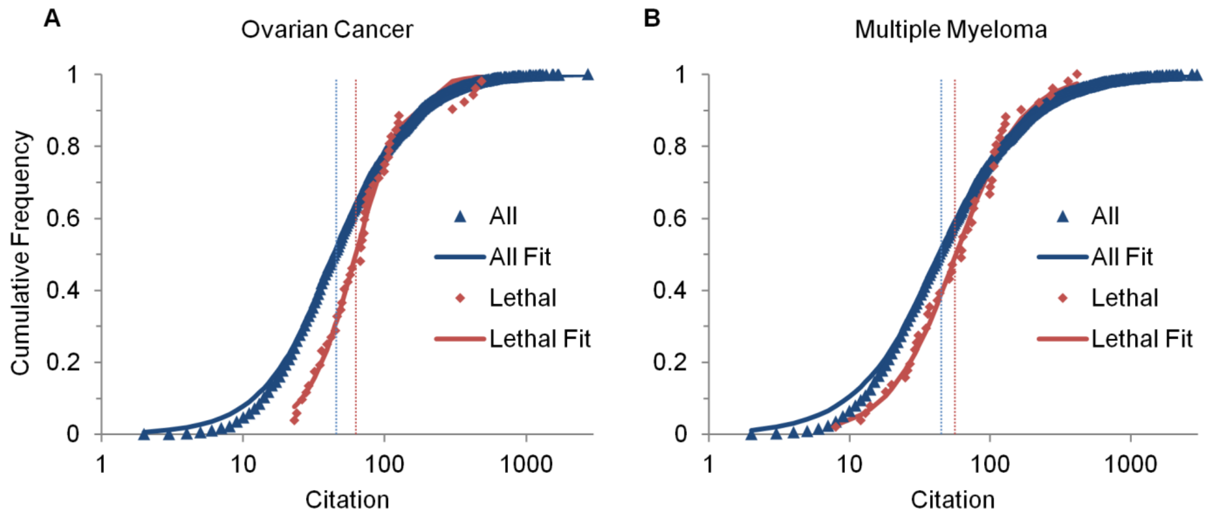

**Figure S4. Destroying neighboring hubs increases network damage.** (A) Scale-free networks were randomly generated using the Barabási–Albert model with 1000 nodes. The two highest-degree nodes were removed to assess if the proximity of the destroyed hubs affects the giant component reduction. The simulation was repeated 10,000 times. Examples of this method applied to networks with hub distance of 1 and 5 are shown. (B) The mean giant component reduction observed in all networks with a given hub distance is shown. Error bars represent standard error of the mean. In networks where the two destroyed hubs were nearer to each other, the giant component reduction was higher than farther hub pairs. The overall trend shows a decrease in giant component reduction with higher network distance between destroyed hubs.

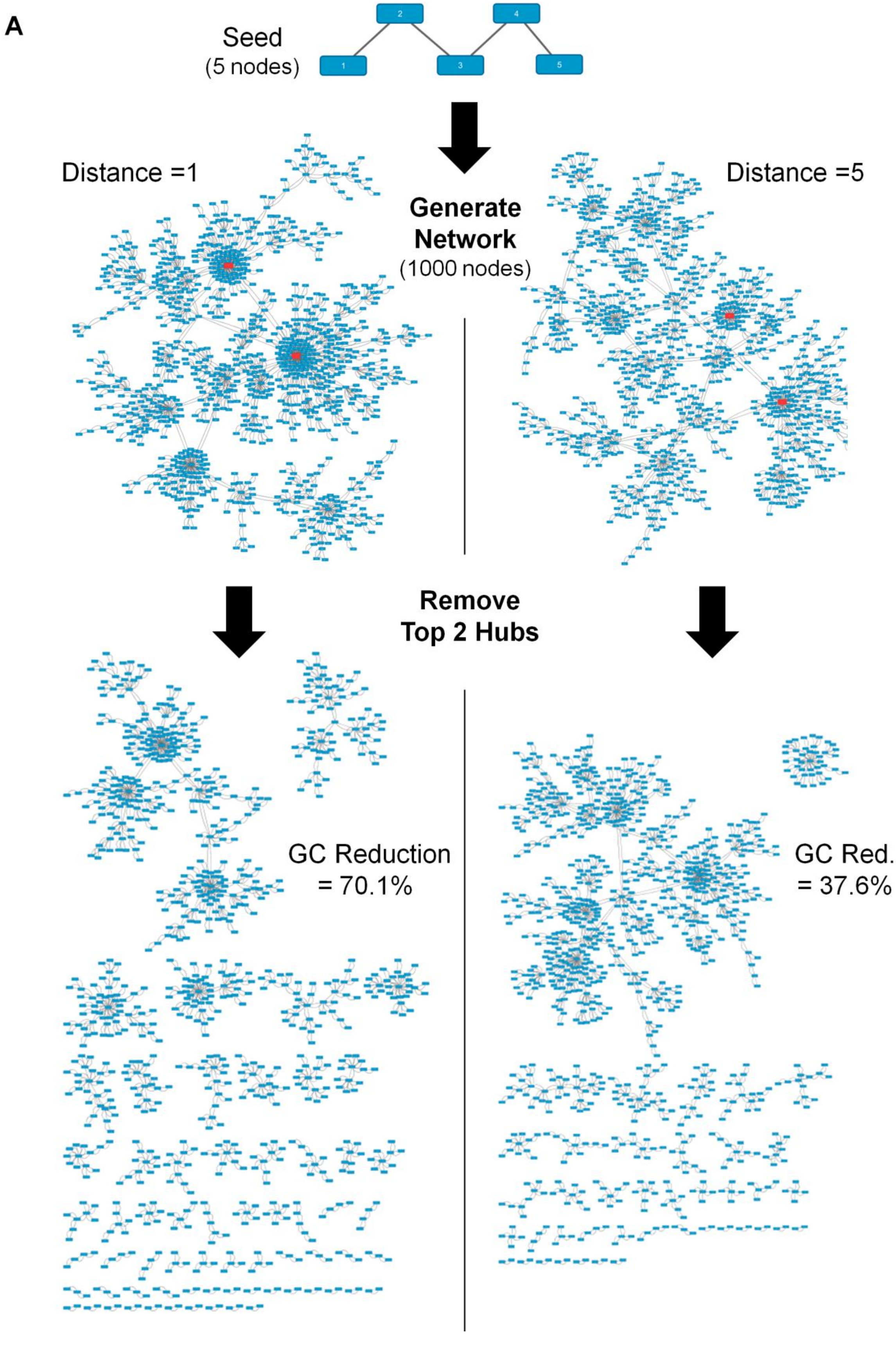
A
Seed
(5 nodes)
1
2
3
4
5
Distance =1
Distance =5
Generate
Network
(1000 nodes)
Remove
Top 2 Hubs
GC Reduction
= 70.1%
GC Red.
= 37.6%

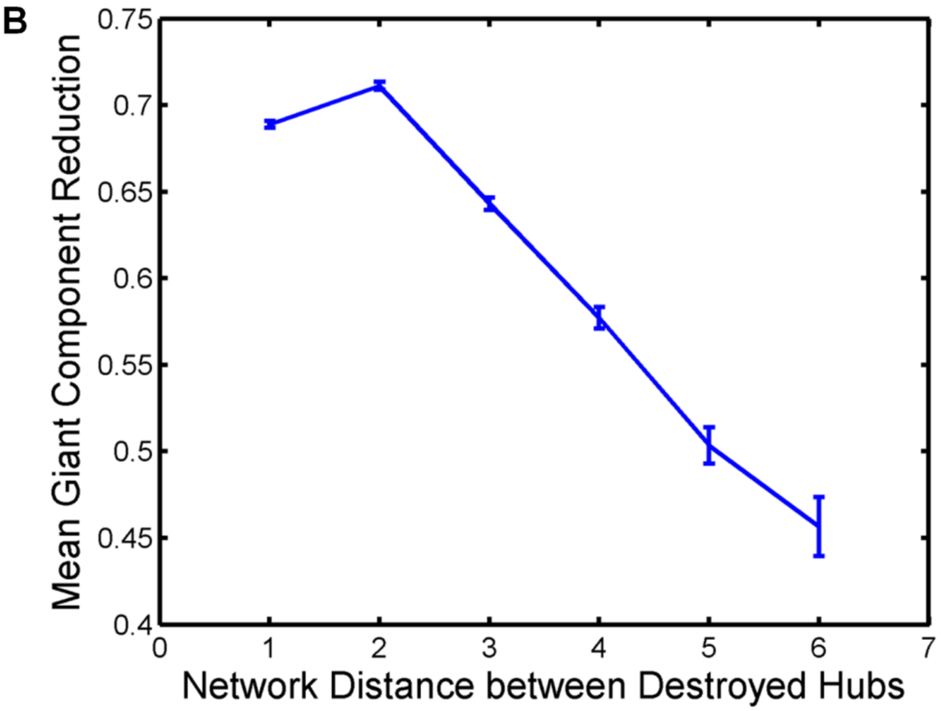
B
Mean Giant Component Reduction
0.75
0.7
0.65
0.6
0.55
0.5
0.45
0.4
0
1
2
3
4
5
6
7
Network Distance between Destroyed Hubs

**Figure S5. Schematic workflow of network analysis.** Genes relevant to 14 cancer indications were extracted from the KEGG PATHWAY database. The BioGRID protein interaction database was searched for interactions involving at least one of a cancer's relevant proteins. This constructed network was then visualized in Cytoscape. Analysis of the network's degree distribution and robustness to node removals was conducted in MATLAB.

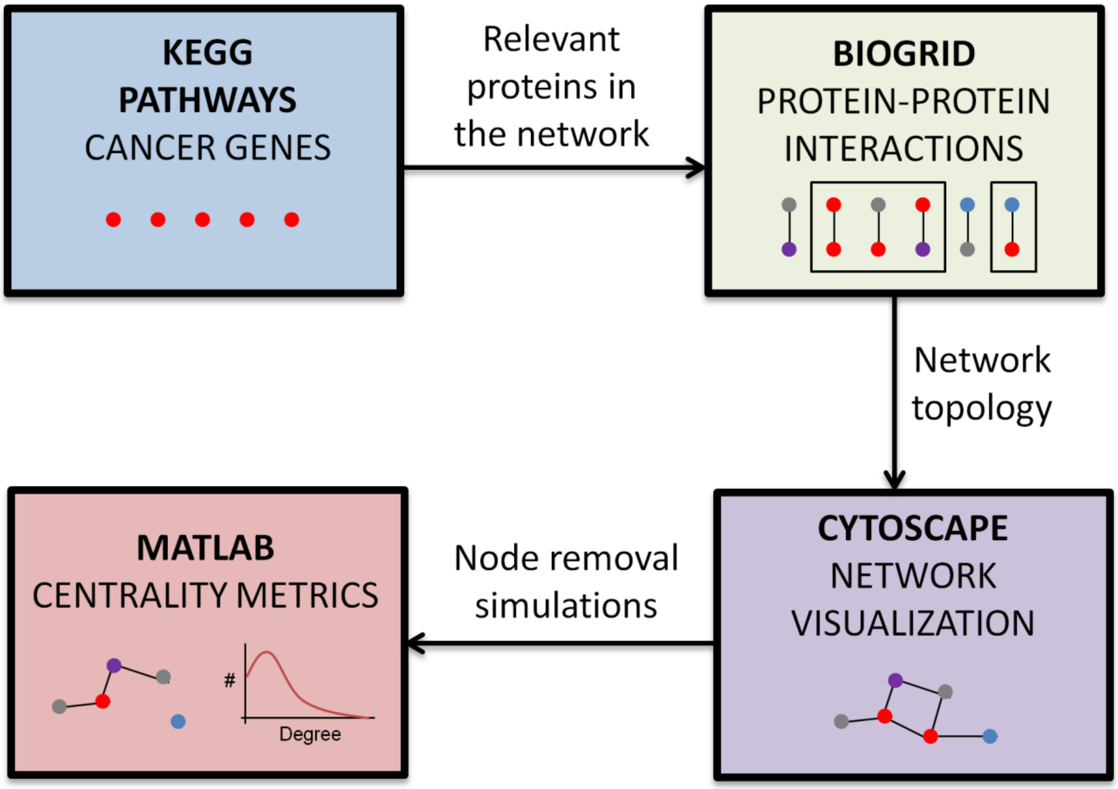
